# Hippocampal connectivity patterns echo macroscale cortical evolution in the primate brain

**DOI:** 10.1101/2023.09.08.556859

**Authors:** Nicole Eichert, Jordan DeKraker, Amy F.D. Howard, Istvan N. Huszar, Silei Zhu, Jérôme Sallet, Karla L. Miller, Rogier B. Mars, Saad Jbabdi, Boris C. Bernhardt

**Author notes:** Lead Contact: Dr Nicole Eichert.

## Abstract

The hippocampus is involved in numerous cognitive functions, some of which have uniquely human aspects, such as autobiographical memory. Hippocampal anatomy, however, is typically considered conserved across primates and its evolutionary diversification is rarely studied. Comparing hippocampal structure and function is, therefore, critical for understanding human brain architecture. Here, we developed a novel comparative framework to study the hippocampus across species characterising its geometry, microstructure, and functional network embedding. In humans and macaques, we generated a new comparative space that represents the hippocampus as an unfolded surface, which respects its sheet-like anatomy. We mapped histological and MRI-derived markers of microstructure to the hippocampal surface and integrated it with low-dimensional embedding of resting-state MRI connectivity data. Our results demonstrate that the micro– and macro-structural organisation of the hippocampus are overall conserved in both species, showing consistent anterior-posterior and subfield-to-subfield differentiation. Furthermore, while hippocampal functional organisation also follows anterior-posterior trends in both species, hippocampal functional connectivity markedly reflected evolutionary reconfiguration of transmodal networks, in particular the default-mode network. Specifically, the inferior parietal lobe in the macaque mirrors an incomplete integration of the default mode network in non-human primates. By combining fine-grained anatomical investigation with large-scale functional imaging, we showed that microstructurally preserved regions like the hippocampus may still undergo functional reconfiguration, due to their embedding in higher-order association networks.

**Summary:** While the hippocampus is key for uniquely human cognitive abilities, it is also a phylogenetically old cortex and paradoxically considered evolutionarily preserved. Here, we introduce a comparative framework to quantify preservation and reconfiguration of hippocampal organisation in primate evolution, by analysing the hippocampus as an unfolded cortical surface that is geometrically matched across species. Our findings revealed an overall conservation of hippocampal macro– and micro-structure, showing anterior-posterior and, perpendicularly, subfield-related organisational axes in both humans and macaques. However, while functional organisation in both species also followed an anterior-posterior axis, the latter showed a marked evolutionary reconfiguration, which mirrors a rudimentary integration of the default-mode-network in non-human primates. Our findings suggest that microstructurally preserved regions like the hippocampus may still undergo functional reconfiguration in primate evolution, due to their embedding in heteromodal association networks.

## Introduction

The hippocampus is one of the most extensively studied parts of the brain given its central role in brain function during health and disease^1^. Indeed, the hippocampus is key to numerous cognitive and affective processes, associated with multiple brain networks, and a model system to examine how neural structure and function covary in space ^2–4^. Importantly, the hippocampus is markedly affected across a range of common and detrimental brain conditions, including neurodegenerative disorders ^5,6^, drug-resistant epilepsy ^7,8^, as well as psychiatric conditions ^9,10^. The hippocampal grey matter consists of archicortex, a phylogenetically old type of cortex, which is considered conserved across mammals^11^. This evolutionary conservation is the basis for cross-species translational frameworks and we have gained a deep understanding for hippocampal anatomy and function from model species, such as non-human primates ^12,13^. It appears to be contradictory, however, that the hippocampus supports many functions that are especially developed in humans for example autobiographical memory^14^, future thinking^15^, and perception of self ^16^. This apparent paradox can be resolved by two potential explanations. Firstly, given that evolutionary diversification of hippocampal anatomy has been rarely studied, it is possible that species differences in hippocampal structure were so far left unnoticed (but see ^17^). Novel quantitative frameworks are required to address this comparison that go beyond measuring regional volumes. Or, secondly, the functional and connectional embedding of the hippocampus within the rest of the brain differs across species and is fundamentally reconfigured since the last common ancestor to humans and monkeys. Species-specific specialisations in subcortical structures such as the striatum ^18,19^, or the amygdala^20^ support the second hypothesis.

The functional embedding of the multiple subdivisions of the hippocampus with the rest of the brain is diverse. On the one hand, it is directly connected to limbic and paralimbic structures, such as amygdala and cingulate cortex, which host some of the most preserved circuits of the brain^21^. On the other hand, it is closely linked to heteromodal regions in neocortex including dorsal-lateral prefrontal and parietal cortex and it forms part of the classical default-mode-network (DMN)^22–24^. The definition of the DMN across the primate lineage, however, is a matter of debate and it has been suggested that it forms two sub-networks in non-human primates^25^. Any evolutionary reconfiguration of the hippocampus is likely related to its functional embedding with the DMN^25,26^, but quantitative evidence for this theory is sparse^27^. Mapping species differences in hippocampal anatomy is, therefore, critical for understanding the origins of human cognition, and also to understand the limitations of model species for neuroscience for studying human disorders such as schizophrenia^28^.

Here, we set out to test these hypotheses in a comparative study between humans and macaques. We devised a novel approach to interrogate cross-species differences in hippocampal anatomy, microstructure, and functional organisation in a common reference frame^29^. We capitalise on recent computational approaches to analytically unfold the hippocampal formation, and to derive a surface-based coordinate system^30^. The topological framework maps the hippocampus intrinsic long (anterior-posterior) and short (proximal-distal) axes, thus respecting the sheet-like anatomy of the hippocampus. Representing the cortex in a surface-based coordinate system has previously proven to advance efforts in brain mapping^31^.

Leveraging this novel common space, we aimed to characterise the sub-regional microstructural organisation of the hippocampus. This work represents the first integration of microstructural features derived from a recently developed multimodal and multiscale macaque atlas including whole-brain post-mortem histological information ^32^ with microstructural features of the human hippocampus ^33^. Then, based on the detailed analytical unfolding of hippocampal anatomy and microstructure we characterised the spatial axes of hippocampal function and its embedding within macroscale functional systems. This then allowed us to test whether spatial axes of hippocampal function underwent a diversification in humans relative to macaques, and whether this diversification coincided with the reconfiguration of cortex-wide functional systems.

## RESULTS

### Hippocampal microstructure is conserved

We adapted a recently developed analytical method for unfolding the hippocampus (hippunfold, ^30^), to the macaque brain, as demonstrated on a template MRI scan based on 10 *ex–vivo* macaques^34^ (**Figure 1A**) for comparison to the human. The 3D surface reconstruction revealed the characteristic seahorse-shape (**Figure 1B**). The unfolded flatmap space is defined based on intrinsic coordinates of the hippocampus ranging from posterior to anterior (from tail to body and head) and from distal to proximal (*i.e.,* from the dentate gyrus to Cornu Ammonis, CA, and subiculum subfields). All hippocampal flatmaps in the remainder of the paper are shown according to this orientation. The hippocampal surface space reconstructed with hippunfold matched geometrically equivalent points across macaque and human. This is demonstrated in **Figure 1C**, which displays geometric indices of the macaque hippocampus next to those from the human.

**Figure 1.**
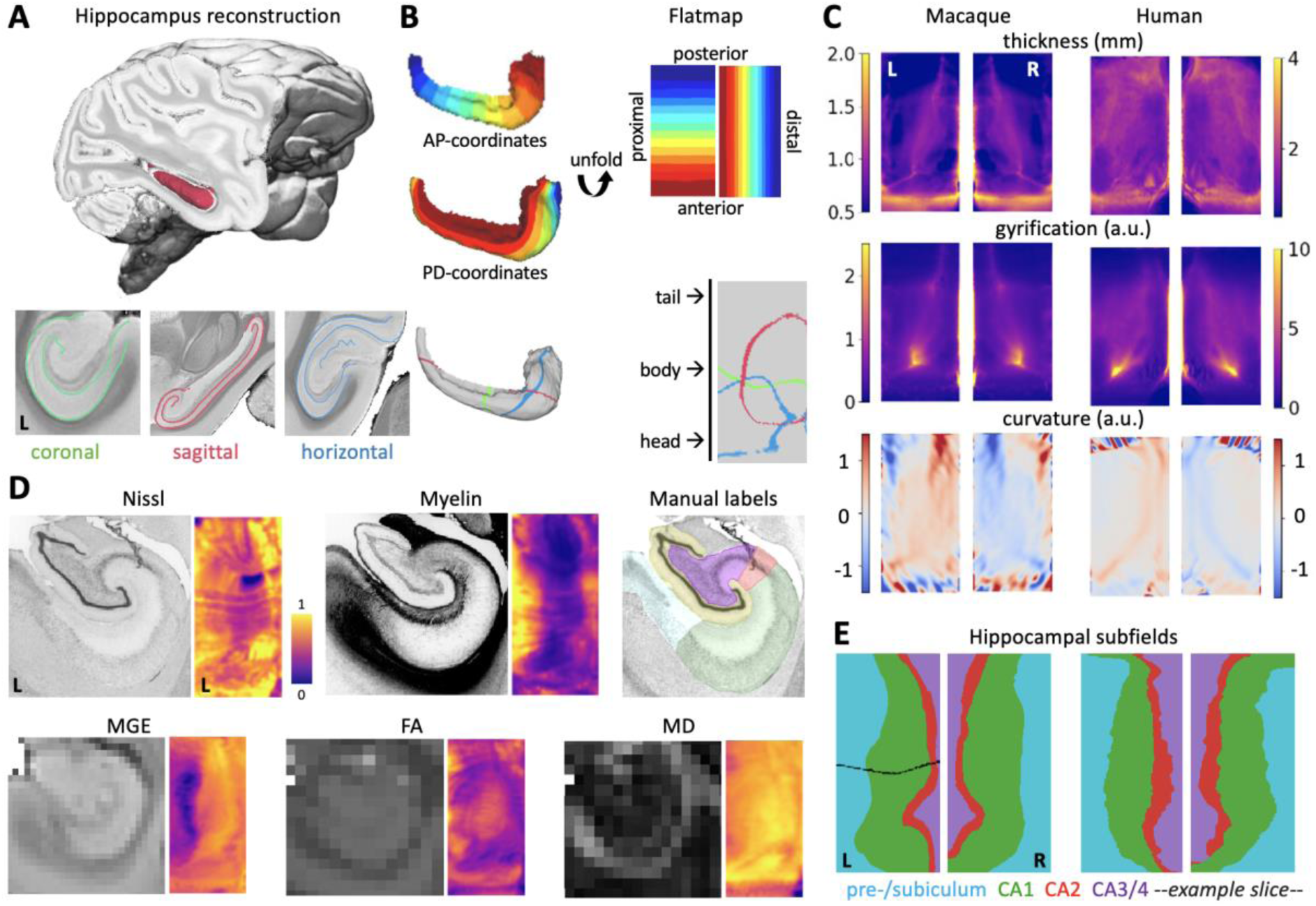
The hippocampal microstructure is conserved. **A**: The hippocampal surface reconstructed in a template MRI scan. **B**: Left – Hippocampal surface as 3D reconstruction and as 2D flatmap. Right – Shown are the intrinsic hippocampal coordinates at the top and the intersections with the planes in A at the bottom. **C**: Geometric indices in hippocampal flatmap space for macaque and human. **D**: Histological and MRI metrics from BigMac. For each modality we show an example coronal section and the whole hippocampus mapped to the flatmap. **E**: Macaque (BigMac) hippocampal subfields and human (BigBrain) subfields. Black line: Intersection with the example histology slice shown in D.

To investigate the hippocampal microstructure in the macaque, we reconstructed the hippocampal surface in a single macaque scan from the BigMac dataset, an open resource combining multi-contrast and ultra-high-resolution microscopy and MRI in a single macaque brain^35^ (**Figure 1D**). We mapped histological measures (Cresyl violet stain for Nissl bodies and Gallyas Silver stain for myelin), manual labels of hippocampal subfields and three MRI metrics (fractional anisotropy [FA], mean diffusivity [MD], multi-gradient-echo intensity [MGE]) to the hippocampal surface. The microstructural mappings demonstrated that hippocampal microstructure varies primarily along the proximal-distal axis and the characteristic spiral configuration of subfields is represented as a sequence of vertical subfields in the flatmap (**Figure 1E**). These variations follow common patterns, for example, the pre-/subiculum has higher intensity in the Gallyas stain and lower MGE intensities compared to CA1, reflecting the higher amount of axons relative to cell bodies, in line with previous reports in the human ^30,36^. We compared the macaque hippocampal map to that from the human BigBrain^33^ and the overall pattern was highly similar (similarity metric of 0.95 and 0.93 for left and right hemisphere, **Supplemental Information, Figure S1G**), despite subtle differences in relative extent of hippocampal subfields were observed. For example, CA2 and CA3/4 are relatively expanded in humans (**Supplemental Information, Figure S1H**). For validation, we repeated hippocampal mapping of the MGE contrast in two additional *ex-vivo* scanned macaque brains, and observed consistent results (**Supplementary Figure S1E**).

Taken together, our hippocampal unfolding and microstructural mapping demonstrated that anatomy and microstructure are overall preserved in both humans and non-human primates, despite subtle changes in global shape and subfield proportions.

### The functional embedding of the hippocampus is diverse

We continued to study the functional anatomy of the hippocampus using resting-state functional MRI (rs-fMRI) data from 10 adult individuals in both species. Rs-fMRI has been widely used to investigate the intrinsic functional brain organisation in both species^37^, and the comparability of functional networks between awake and lightly anaesthetised states has been firmly established ^38–40^. Image pre-processing and analysis were equivalent in both species and the macaque fMRI data are of higher tSNR than most previously used datasets ^41^, altogether suggesting that the data in both species allow for a quantitative comparison.

Application of non-linear dimensionality reduction techniques^42^ to hippocampus-to-cortex connectivity matrices provided spatial maps of connectivity variations, also referred to as *Connectivity Gradients*^22,43,44^, of the hippocampus (**Figure 2A**). The first component revealed a pronounced anterior-posterior axis in both species (**Figure 2B**). Later components (2^nd^ in the human and 6^th^ in the macaque) show further differentiation along the proximal-distal axis (**Supplemental Information, Figure S2A**).

**Figure 2.**
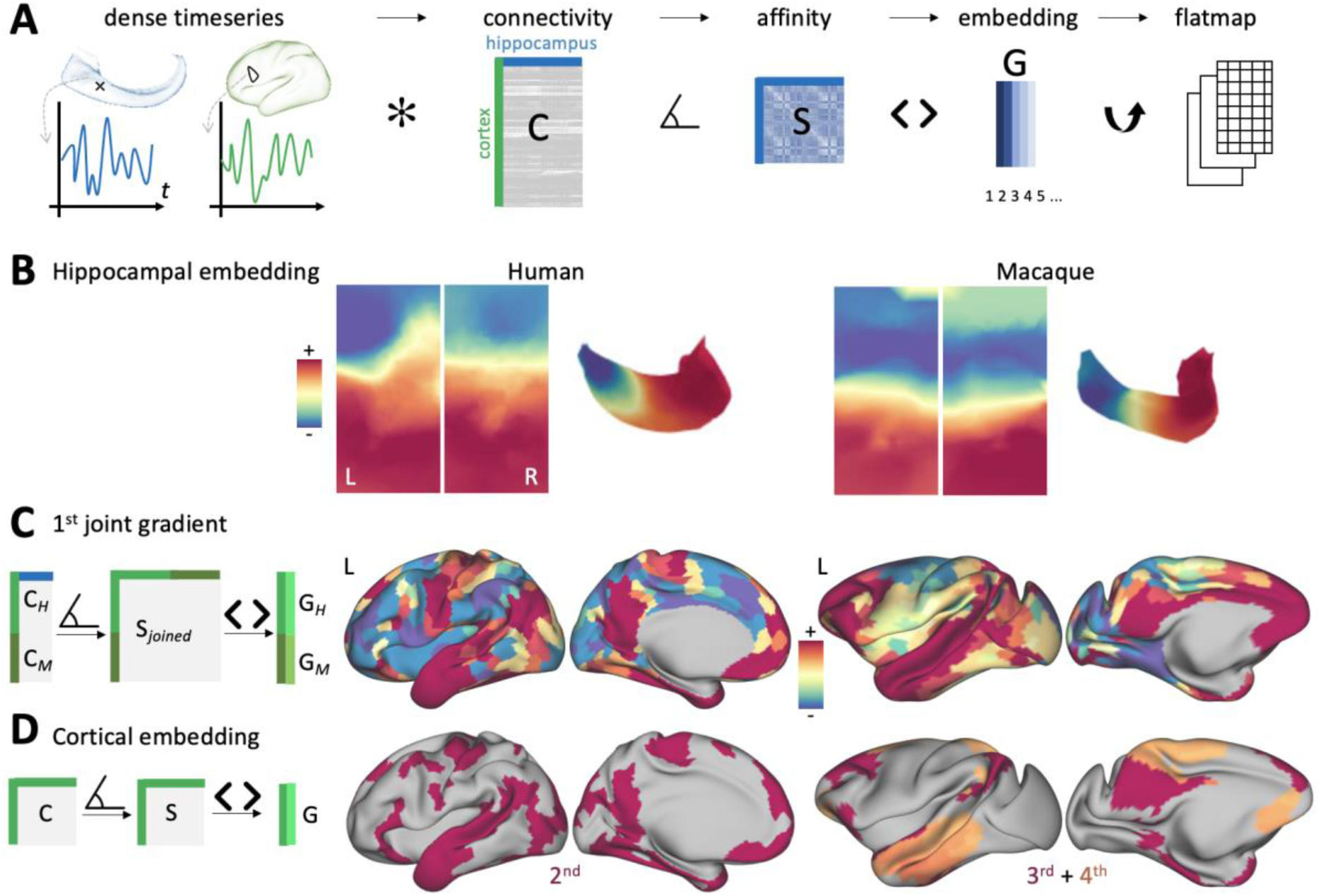
The functional embedding of the hippocampus is diverse. **A**: Workflow to perform hippocampal gradient embedding. A dense connectivity matrix (C) is constructed from the hippocampal and cortical resting-state data. After applying a similarity operation, the square matrix (S) is decomposed into gradient maps (G). **B**: First hippocampal gradient in humans and macaques shown as flatmap and on the left hippocampal surface reconstruction. **C**: Left – A joint cross-species cortico-hippocampal gradient (G) was obtained by concatenating the human cortico-hippocampal connectivity matrix (C_H_) with that of the macaque (C_M_) followed by an affinity operation (S) and gradient embedding (G). Right – 1^st^ joint cross-species gradient in human (G_H_) and macaque (G_M_). **D**: Left – In each species separately, cortico-cortical gradients were computed. Right – Thresholded maps of cortical gradients that match the joint cross-species gradient.

Next, we characterised the functional connectivity of the hippocampus with the rest of the cortex (**Figure 2C**). To this end, we performed simultaneous decomposition of cortex-to-hippocampus connectivity data in both species. This ‘joint connectivity gradient’ mapping approach provided us with homologous spatial maps of connectivity variation across the cortex. One apex of the 1^st^ component (warm colours in Figure 2C) recovered a network of homologous areas in both species including posterior cingulate/precuneus, lateral temporal lobe, supramarginal gyrus, ventro-medial and dorso-lateral prefrontal cortex. The regions within this network are all highly connected to the hippocampus (**Supplemental Information, Figure S3A**). In both species, hippocampal connectivity was also observed to inferior temporal gyrus and occipital lobe, but to a lower extent. Some species differences in this network, however, were found as well: For example, in macaques, the supramarginal gyrus and somatosensory cortex are less differentiated, compared to the human. Overall, however, our analyses uncovered a homologous brain network in both species, defined by similar connectivity profiles to the hippocampus.

In humans, the resulting network from our hippocampal analysis is reminiscent of a default mode network (DMN). This finding is in line with our expectation, as the hippocampus is a well-established node of the human DMN^45^. The macaque network that emerged from the same analysis, can be thought to represent the macaque homologue of the DMN. However, previous definitions of the macaque DMN using, for example, ICA suggested that the macaque DMN consists of two sub-networks^25^ or only a partial network^26,46^. To demonstrate that the homologous macaque DMN, as we defined it using hippocampal connectivity, in fact comprises two distinct cortical networks, we conducted additional analyses.

First, we performed dimensionality reduction of cortico-cortical connectivity matrices in both species and compared these to the hippocampal network from the joint connectivity gradient analysis above. All cortical networks from both species are shown in the **Supplemental Information, Figure S2B**. As expected in humans, one cortical network map matched the joint gradient best (regularised regression analysis, coefficient = 0.76, r^2^ = 0.55, *p_corrected_* < 0.001). The best fit for the joint gradient in the macaque was found with the 4^th^ cortical embedding map (coefficient = 0.46, r^2^ = 0.16, *p_corrected_*< 0.001), closely followed by the 3^rd^ (coefficient = 0.21, r^2^ = 0.06, *p_corrected_* < 0.05). Successive Dice overlap analysis confirmed that a combination of map 3 and 4 in the macaque matches the joint gradient best (Dice = 0.54, at threshold 75%). This analysis confirmed that the macaque homologue of the DMN as defined by hippocampal embedding is best explained by a combination of two distinct cortical networks: The 4^th^ cortical embedding map recovers the lateral temporal and medial frontal DMN nodes, whilst the 3^rd^ gradient recovers the inferior parietal, the dorso-lateral prefrontal and the precuneus DMN nodes (**Figure 2D**). Precisely these two subnetworks have previously been suggested to form the DMN in non-human primates ^25^.

To extend the species comparison to a whole-brain level, we finally computed a vertex-wise homology index between species based on the connectivity profiles with the hippocampus ^47,48^. This analysis confirmed that somatomotor and limbic networks are most conserved across species, whilst higher-order networks such as the DMN and multiple-demand network, as defined by the discrete Yeo human network parcellation^49^, are more strongly reconfigured (**Supplemental Information, Figure S3C**).

Our functional analysis showed that the hippocampus is embedded with homologous large-scale functional networks in both species with strongest involvement of the DMN. Whilst the DMN-homologue in the macaque forms two distinct sub-networks on the cortical level, the full macaque DMN functionally interacts with the hippocampus. Taken together with the histological results above we showed that the ‘short’ proximal-distal hippocampal axis captures microstructural variations of the hippocampus, whilst the orthogonal ‘long’ anterior-posterior characterises variations in its functional organisation.

### Cortical embedding of the hippocampus reflects evolutionary reorganisation

Finally, we studied how the functional topography of the hippocampus is reflected in macroscale cortical networks and explicitly investigated how the two established hippocampal axes map onto the cortex. We used dual-regression ^50,51^ to determine the individual contribution of the two orthogonal hippocampal gradients to cortical connectivity (**Figure 3A**).

**Figure 3.**
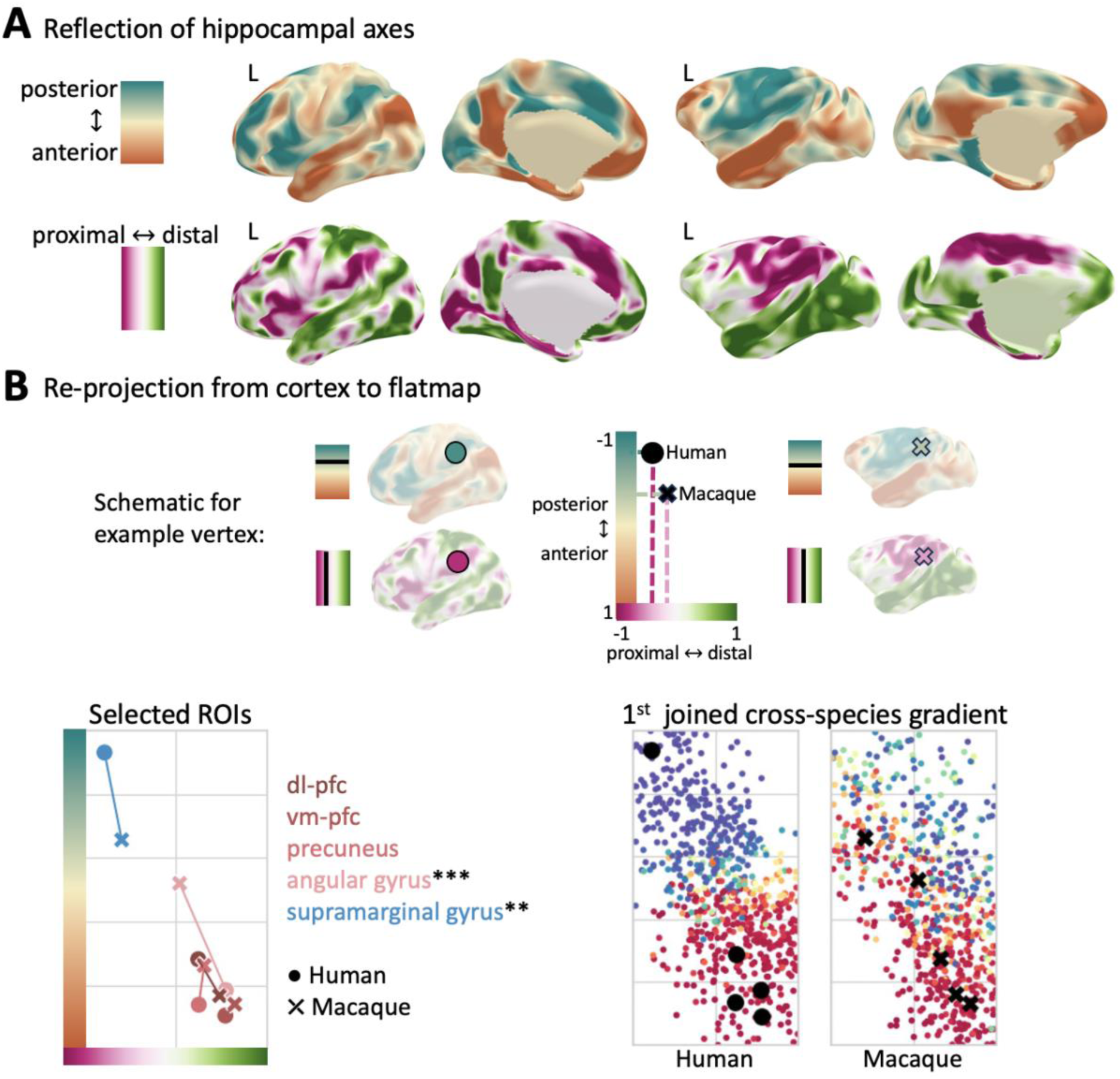
Cortical embedding of the hippocampus reflects evolutionary reorganisation. **A**: Cortical reflection of the hippocampal axes derived using dual regression. The colours in the cortical maps (right) represent differential connectivity to different parts of the hippocampus (left). Colormaps are matched for both species (10% – 90%) **B**: Top – Schematic diagram to demonstrate how an example cortical vertex is remapped to the 2D flatmap space based on its connectivity in A. The aspect ratio of the coordinate system reflects that of the hippocampal space. Bottom left – Five brain areas in both species mapped onto the 2D space. We tested the difference between human and macaque using two-sample Kolmogorov-Smirnov test (\*\*\**p*<0.001, \*\**p*<0.01, *n*=10). Bottom right – 1^st^ joint cross species gradient (Figure 2C) represented in a 2D space. Black data points are the selected brain areas shown in the right panel.

The cortical reflection of the long-axis gradient shows that the anterior part of the hippocampus notably mediates connectivity with the DMN nodes in both species (warm colours in Figure 3A: angular gyrus, middle temporal lobe, posterior cingulate/precuneus, dorso-lateral frontal and anterior medial as well as orbital frontal cortex). Preferential connectivity with the posterior hippocampus, however, displays a pattern previously described as multiple-demands-network^52^. A notable species difference, however, is observed in the inferior parietal lobe, which shows a clear dissociation along the anterior-posterior axis in the human, but little differentiation in the macaque. Connectivity maps of spatially distinct hippocampal segments confirm these patterns (**Supplemental Information, Figure 3B**).

The cortical reflection of the proximal-distal axis reveals a more distributed pattern. Connectivity to DMN nodes, but also to higher-order visual areas and ventral premotor cortex are mediated by the distal part of the hippocampus. Again, the inferior parietal lobe shows a clear distinction within this axis in the human, but not in the macaque.

The differential connectivity along the two hippocampal axes allows us to map cortical brain areas back into a two-dimensional coordinate system, which in turn represents the intrinsic hippocampal space (**Figure 3B**). We leveraged this visualisation to map selected brain areas: Four nodes of the DMN (*ROIs with warm colours*) and the supramarginal gyrus (*ROIs with cold colours*). This visualisation highlights that the DMN nodes from both species map onto similar locations in the shared space. The two nodes of the inferior parietal lobe (angular and supramarginal gyrus) exhibit a strong dissociation in humans falling onto opposite corners of this space. In the macaque, however, both inferior parietal nodes show much reduced differentiation along both axes. We further reprojected the joint cross-species gradient from above (Figure 2C) to this space, demonstrating that connectivity with this hippocampal network is mediated by anterior and distal parts of the hippocampus in both species.

Taken together, these results demonstrate that the DMN nodes exhibit differentiated connectivity with the hippocampus reflecting the intrinsic topography. The macaque inferior parietal lobe reflects incomplete integration of the DMN in the non-human primate.

## Discussion

Studying the hippocampus, a key interface of paralimbic and heteromodal association systems, provides important insights to revise our understanding of primate evolution and conservation^11^. The present paper devised a novel comparative framework to study evolutionary reconfiguration of hippocampal microstructure, anatomy, and integration into large-scale systems. To this end, we successfully unfolded the non-human primate hippocampus for the first time using a recently developed analytical tool^30^. The hippocampal 2D surface is geometrically matched across species and, therefore, represents a novel common space for comparative analyses^29^. First of all, we found that the microstructural blueprint of the hippocampus, its principal functional axes in anterior-posterior direction, and cortical network embeddings are overall conserved across species. However, hippocampal embeddings to macroscale functional also reflected evolutionary innovation, in particular the default-mode network (DMN). Specifically, the inferior parietal lobe in the macaque mirrors an incomplete integration of the DMN in non-human primates. Our findings thus suggest that the human DMN has expanded and further integrated in the human lineage, harnessing the hippocampal microcircuit in a uniquely human way. Altogether these adaptations form the basis of specialised human brain function spanning a wide range of cognitive functions. While the hippocampus’ structural organisation is largely conserved, its functional connectivity has evolved, demonstrating that even structurally preserved regions like the hippocampus can undergo functional adaptations due to their connections with higher-order networks.

We leveraged two unique ultra-high-resolution multiscale databases from the macaque and human for a cross-scales and cross-species comparison. This novel combination of resources allowed us to demonstrate that the hippocampal macro– and microstructure are overall conserved across species, a long-standing evolutionary hypothesis that was lacking spatially resolved quantification to date. The histological mapping further functioned as a microstructural validation of our hippocampal surface and flatmap, confirming that it represents a comparative space across primates. Our findings overall show that both species have a comparable long axis, comprising head, body and tail regions that are characterised by a canonical sequence of subfields. This demonstrates that microstructural patterns and gradients in the hippocampus are conserved across species, suggesting that the basic microcircuit and hippocampal computation remained largely unchanged over evolutionary time. Despite the pronounced similarities in subfield mapping across species, however, we also showed that our framework can detect nuanced differences. For example, our findings revealed a relative expansion of the CA2 subfield in the human, an effect that may be compatible with the hypothesised functional specialisation of CA2 for social memory ^53^ and territorial behaviour^54^, underpinned by the subfields distinct cytoarchitecture^52^ and genetic profile^55^. While findings on potential cross-species microstructural differences require further validation in a larger histological sample to discern inter-species from inter-individual differences, the methodology we introduced here offers a scalable framework to allow for microstructural comparisons across humans and non-human primates. Beyond their value for fundamental neuroscientific inquiry, as carried out in the current study, this approach may also be beneficial when translating findings between animal models and patient groups in a preclinical/clinical context. Methods to automatize hippocampal subfield segmentation ^30,56,57^ and to enhance cross-modal and inter-individual registrations^30^ are already well established in humans, and current efforts to adapt them to non-human primate brains will facilitate that process. In this context, open data sharing projects such as the BigMac dataset^32^ used in this study, but also initiatives such as PRIME-DE^58^ may make an invaluable contribution, as they will allow for the aggregation of a diverse set of data in non-human primates, and their dissemination to a wide range of researchers.

Building up on the microstructural analysis, our resting-state fMRI results demonstrated that the functional differentiation along the hippocampal long axis is also largely preserved across species. The cortical embedding of the hippocampus recovered a homologous default-mode network in both species comprising dorso-lateral frontal, inferior-parietal, anterior-temporal, as well as fronto– and posterior-medial regions. Importantly, this homologous DMN was defined based on joint embedding of hippocampal connectivity and did not require the specification of homologous cortical regions-of–interest. Because of the data-driven nature, anchored only by the hippocampus, our approach is readily applicable to all mammalian species, and represents an important further development to previous remapping approaches ^47,48^. General limitations of cross-species comparisons using resting-state fMRI were mitigated by using a macaque dataset with high temporal signal to noise, long scanning time, light anaesthesia and by using the equivalent HCP preprocessing pipeline in both species. Despite the overall similarity of functional networks, we also observed notable species differences in functional connectivity. In particular, we showed that the DMN, known to act as a major integrated network in humans ^24,45,59^, constitutes two subcomponents in the macaque, when studied on the cortical level only. Various previous definitions of the non-human primate DMN homologue referred only to one of the two subnetworks^46^ or described them as two distinct subnetworks ^25^. Our analysis suggests, however, that all conventional DMN nodes are present as precursors in the macaque. The critical species difference we propose is that macaques exhibit incomplete integration of the DMN, whilst humans have a fully integrated network. The increase in network integration and connectivity in the human lineage mirrors the increase in DMN integration during typical human brain development, seen both functionally ^60,61^ and structurally ^62–64^. Our findings thus provide new evidence to support the hypothesis that the computational landscape of the mature adult brain is fundamentally linked to the development of long-range connections and the development of the DMN. The cortical embedding of the two hippocampal axes revealed that the inferior parietal lobe in the macaque specifically reflects this incomplete network integration. Our results suggest that parietal connectivity to the temporal lobe and posterior medial cortex are reduced in macaques, which is in line with previous comparative literature on structural ^65,66^ and functional connectivity ^37,47^. In addition to changes in connectivity, the parietal lobe is one of the hotspots of cortical expansion, which is supported by uniquely human genetics^67^. These adaptations support the specialised role of the parietal lobe in social cognition^68^ and theory of mind^69^ in humans. Our findings, therefore, reconcile and extend previous controversies about the evolution of the DMN ^26^ and the functional significance of DMN integration for mature human brain function.

A wealth of data from multiple species and modalities suggests that hippocampal organisation can be conceptualised along two quasi-orthogonal axes ^22,52^. The spatial gradients we proposed in this paper therefore serve as a meaningful coordinate system to study preservation and innovation in brain evolution. The ‘short’ proximal-distal axis captures microstructural variations traditionally described as discrete subfields^52,70,71^. These structural variations have been well established in a range of mammalian species and they are the neural substrate for a specialised circuitry that forms the basis of hippocampal function^72^. Indeed, our histological analysis demonstrated that the subregional organisation and, therefore, the canonical hippocampal microcircuit along the proximal-distal axis is largely conserved across the primate lineage. The spatial topology of the hippocampus, however, is also characterised by variations along the ‘long’ anterior-posterior axis^4^, which recapitulates the segmental head-body-tail anatomical arrangement. Notably, spatial patterns in gene expression^73^, receptorarchitecture^74^ and hippocampal function ^75^ have been shown to closely follow this long axis in the human. Our results provide one of the first empirical demonstrations of a long-axis functional differentiation in non-human primates (see also ^76^) and demonstrate direct correspondence to the human. The diverse brain-wide connectivity of the hippocampus is thought to mirror the functional substrates of hippocampal involvement in various cognitive and behavioural domains ^77^. In line with this theory, our analysis demonstrated that the functional connectivity of the hippocampus is diverse and varies along both intrinsic axes. Overlapping and covarying axes of microstructure, connectivity, and molecular profiles together can possibly explain how the hippocampus can be engaged in many different brain functions ^3^ ranging from relational memory across spatial, episodic and semantic contexts ^78–81^ to other domains including emotional reactivity ^22,82^ and stress^83^. Whilst we found that the fundamental microstructure of the hippocampus is overall preserved, species differences in the functional embedding of the hippocampus are possibly mirrored in nuanced species differences on the level of gene expression or receptorarchitecture ^84^.

Taken together, our work provides quantitative evidence for long-standing theories of brain evolution. The multisynaptic pathway of the hippocampus likely emerged in early mammals to promote survival in complex environments via spatial navigation and pattern separation ^78,85^. According to existing theories, rodents leverage this basic computation for integration of proximal and spatial cues ^86^, whilst primates repurpose the circuit to integrate visual and abstract cues^79^ or values^87^. This integration relies on extended connectivity with other brain regions and eventually gives rise to higher-order cognitive abilities, such as episodic memory^88^ and social cognition^89^. Hippocampal function therefore essentially co-evolved with heteromodal systems such as the DMN and at the same time maintained its capacity to integrate sensory processing streams ^90,91^. The preservation of a successful microcircuit and simultaneous ‘gain of function’ by virtue of specialised cortical connectivity makes the hippocampus a prototypical region to understand human brain evolution. This interpretation of our findings fits in well with the growing body of literature of evolution, demonstrating that human brain evolution is a nuanced process that goes well beyond brain expansion. On the level of cortical brain areas, cross-species work across the mammalian family has demonstrated that multiple forms of adaptations can overlap and interact, such as relative size of cortical fields and changes in connectivity ^29,92^. This concept extends to the brain network level as previously demonstrated for the language system in the brain ^93^. Regional expansions of the cortex, for example, do not explain the changes to long-range white matter tracts across primates^94^. These regional expansions are most pronounced in association cortex and mirror patterns of brain expansion during human development^95^. It has been suggested that these rapid expansions free up potions of cortex that become ‘untethered’ or from early molecular constraints imposed by conserved brain anchor regions. The untethering of these regions to develop more dense connections within and between brain networks^96^. Our results align well with the untethering hypothesis, where the hippocampus forms a conserved anchor for the primate DMN, which increasingly expanded and integrated in humans thus endowing the hippocampus with increased functionality.

In conclusion, we developed a novel comparative framework to study the hippocampus across species. Our in-depth study of the primate brain integrated ultra-high-resolution assessments of hippocampal microstructure with advanced decompositions of its functional network embedding, and demonstrated how conserved brain regions can functionally adapt through interactions with advanced networks. We anticipate this paper to be the starting point for a new generation of comparative studies, to unlock a deeper understanding of the evolution of our own cognitive abilities.

## Acknowledgements

NE is supported by a Sir Henry Wellcome Postdoctoral Fellowship from the Wellcome Trust [222799/Z/21/Z]. JdK is supported by a Natural Sciences and Engineering Research Council of Canada – Postdoctoral Fellowship (NSERC-PDF). AFDH and INH were funded by the Wellcome Trust [WT215573/Z/19/Z]. KLM and the BigMac dataset were funded by the Wellcome Trust [WT202788/Z/16/Z]. SZ was supported by the Chinese Government Scholarship. SJ is supported by a Wellcome Collaborative Award [215573/Z/19/Z] and a Wellcome Senior Research Fellowship [221933/Z/20/Z]. BCB acknowledges research support from the National Science and Engineering Research Council of Canada (NSERC Discovery-1304413), CIHR (FDN-154298, PJT-174995), SickKids Foundation (NI17-039), Helmholtz International BigBrain Analytics and Learning Laboratory (HIBALL), Healthy Brains and Healthy Lives (HBHL), BrainCanada, and the Tier-2 Canada Research Chairs program. The Wellcome Centre for Integrative Neuroimaging is supported by core funding from the Wellcome Trust [203139/Z/16/Z and 203139/A/16/Z]. For the purpose of Open Access, the author has applied a CC BY public copyright licence to any Author Accepted Manuscript version arising from this submission.

## Author contributions

Conceptualization, N.E. and B.C.B.

Methodology, N.E., J. DK., A.F.D.H., I.N.H., S.Z., R.B.M., S.J., B.C.B.

Software, J.D.K., I.N.H.

Resources: J.S., A.F.D.H., K.L.M., R.B.M.

Writing – Original Draft, N.E. and B.C.B.

Writing – Review & Editing, all authors.

## STAR METHODS

### RESOURCE AVAILABILITY

#### Lead Contact

Further information and requests for data and code should be directed to and will be fulfilled by the lead contact, Dr Nicole Eichert

#### Materials Availability

This study did not generate new raw data and only accessed previously acquired and published data.

#### Data and code availability

● HCP data are publicly available at https://www.humanconnectome.org/. The CIVM macaque post-mortem template is publicly available at https://civmvoxport.vm.duke.edu/. Histological and MRI data as well as the slide-to-volume registrations from BigMac are existing and publicly available via the Digital Brain Bank platform (https://open.win.ox.ac.uk/DigitalBrainBank). Additional macaque MRI in-vivo data will be made publicly available as of the date of publication via the PRIME-DRE repository (https://fcon_1000.projects.nitrc.org/indi/indiPRIME.html). The hippocampal segmentation used for unfolding the macaque hippocampus in the template macaque scan will be shared via the Wellcome Centre of Integrative Neuroimaging’s GitLab server as of the date of publication.
● Hippunfold is openly available as a BIDS App at https://github.com/khanlab/hippunfold. All original code generated for this project has been deposited at the Wellcome Centre of Integrative Neuroimaging’s GitLab server (https://git.fmrib.ox.ac.uk/neichert/project_hipmac) as of the date of publication.
● Any additional information required to reanalyse the data reported in this paper is available from the lead contact upon request.

### EXPERIMENTAL MODEL AND STUDY PARTICIPANT DETAILS

#### Human – MRI data

Human MRI data were provided by the Human Connectome Project, WU-Minn Consortium (Principal Investigators: David Van Essen and Kamil Ugurbil; 1U54MH091657) funded by the 16 NIH Institutes and Centers that support the NIH Blueprint for Neuroscience Research; and by the McDonnell Center for Systems Neuroscience at Washington University. We accessed minimally pre-processed structural and resting-state fMRI data in fsLR_32k space of 10 subjects from the 3T 1200 Subjects Data Release^97^ (4 females, mean age 30.4 ± 3.2 years) to match the sample size in the macaque. The subjects were randomly selected.

#### Human – Histological data

Previously established human histological subfield mapping was based on a single post-mortem sample from a 65-year-old male donor, BigBrain^33^. We accessed only the surface map of discrete hippocampal subfields, which are provided via open-access by a previous study ^98^.

#### Macaque – Post-mortem MRI data

A high-resolution (0.15 mm^3^) rhesus macaque *(Macaca mulatta*) structural gradient-echo macaque template based on 10 individuals was accessed from the CIVM Macaque Brain Atlas ^34^. Furthermore, we accessed 7T post-mortem whole brain data from a male adult rhesus macaque, the BigMac Dataset^35^, provided openly accessible by the Oxford Digital Brain Bank (open.win.ox.ac.uk/DigitalBrainBank)^99^. Full details of the BigMac data acquisition and preprocessing are provided in the original publication^35^. Specifically, we accessed the multi-gradient-echo (MGE) high-resolution structural scan (0.3 mm^3^) as well as pre-processed fractional anisotropy and mean diffusivity data from the b=4k diffusion acquisition (0.6 mm^3^). Histological data (see below) from the same individual was also accessed. Post-mortem MGE scans were acquired using the same sequence in two additional macaque brains.

#### Macaque – *In-vivo* MRI data

All macaque in-vivo data were acquired for previous studies ^37,39^ and reanalysed for the purpose of the present paper. Resting-state functional and in-vivo structural MRI data were obtained from 10 rhesus macaques (*Macaca mulatta,* 1 female, mean age at scan 7.2 ± 2.5 years). Details of the scanning protocol and physiological monitoring are described in a previous publication ^37^. In short, the animals were scanned in a 3T scanner under light isoflurane anaesthesia whilst placed in an MRI compatible stereotactic frame or resting on a custom-made mouth mould. In short, BOLD fMRI was acquired for 1600 volumes (∼1 h) with the following parameters: 1.5 mm^3^ spatial resolution, TR = 2280 ms, TE = 30 ms. Structural scans using a T1-weighted MPRAGE sequence were acquired at 0.5 mm^3^ during the same scanning session.

#### Macaque – Histological data

We accessed microscopy data from the BigMac dataset, specifically the Cresyl Violet and the Gallyas Silver stain sections, which had been digitally downsampled to a resolution of 40 μm. We also accessed previously generated registrations for each histology slice to the MGE volume. These registrations were derived using FSL’s TIRL ^100,101^.

#### Ethics Statement

All HCP scanning protocols were approved by the local Institutional Review Board at Washington University in St. Louis. All subjects provided informed consent prior to participating in the study. The donor for the post-mortem BigBrain sample is not personally identifiable and gave written informed consent for the general use of post-mortem tissue used in this study for aims of research and education. The usage of the post-mortem material is covered by a vote of the ethics committee of the medical faculty of the Heinrich Heine University Düsseldorf (#4863). All experimental procedures in macaques were performed in compliance with the United Kingdom Animals (Scientific Procedures) Act of 1986. A Home Office (UK) Project License, obtained after review by the University of Oxford Animal Care and Ethical Review Committee, licensed all procedures. The housing and husbandry followed the guidelines of the European Directive (2010/63/EU) for the care and use of laboratory animals. The 3Rs principles compliance and assessment was conducted by the UK National Centre for 3Rs (NC3Rs).

### METHOD DETAILS

#### Hippunfold

To reconstruct the hippocampal surface, we used a recently developed computational tool, hippunfold v.0.3. Hippunfold requires a tissue segmentation to define unfolded coordinate boundaries of the hippocampal surface. In humans, the segmentation can be derived automatically by hippunfold using a convolutional neural network, but an adaptation for the macaque brain required us to manually segment a set of hippocampus and surrounding structures. We developed the macaque segmentation in a high-resolution (0.15 mm^3^) gradient-echo MRI scan from the CIVM database. The following labels were manually segmented in ITK-SNAP v.3.8.0^102^: The hippocampal grey matter, the dentate gyrus, stratum radiatum, lacunosum and moleculare (SLRM, the ‘hippocampal dark band’), the grey matter of the temporal lobe adjacent to the hippocampus, the uncus, the hippocampal-amygdalar transition area, and indusium griseum (**Supplemental Information, Figure S1A**). We followed the segmentation protocol developed for the human^103^, with minor notable differences: i) the macaque hippocampus was less gyrified and so segmentation of grey matter was simpler, ii) the uncus of the hippocampus was smaller in the macaque, but still showed the same critical termination on the amygdala that allows for unfolding, and iii) the boundary between subiculum and medial temporal lobe neocortex was shifted laterally in the macaque compared to the human, making the macaque hippocampus smaller, to accommodate the darker intensity of the parasubiculum which was visibly shifted laterally in macaques compared to humans.

The segmentation was developed in the left hemisphere, then initialised in the right hemisphere by non-linear registration using ANT’s QuickSyN tool ^104^ and manually corrected in the right hemisphere. Using the segmentations as input, hippunfold was run using default settings, and two surface meshes at a resolution of 419 vertices per hemisphere (‘low-resolution mesh’, **Figure S1B**) and 7262 vertices (‘high-resolution mesh’) were obtained. In addition to the outer, inner, and mid-thickness surface, we obtained in total 6 equivolumetric surfaces across the hippocampal depth. The hippocampal flatmap space is derived by hippunfold by estimating the 2D Laplacian coordinates of the grey matter sheet. An expansion showing the distortions of the surface is provided in **Figure S1C**. Hippunfold automatically computes vertex-wise measures of hippocampal grey matter thickness, gyrification and curvature. Furthermore, the previously labelled human hippocampal subfield labels from BigBrain are provided as a categorical surface map.

For resting-state analysis (see *below*) the hippocampal surfaces from the CIVM template were non-linearly transferred to the Yerkes-19 macaque template space ^105,106^. In the MNI template brain, hippunfold was run using default settings with automated segmentation. For histological mapping in the macaque BigMac brain, we manually corrected the tissue segmentation following nonlinear initialization to fit the individual’s anatomy optimally and then ran hippunfold in the BigMac brain as described above.

#### Subfield drawing and microstructural mapping

Hippocampal subfields were manually labelled in QuPath v.0.2.3 onto the Cresyl Violet stained histological slices (see **Figure 1D** for an example). The cutting angle of the slices relative to the hippocampal long axis was approximately 50° and the slice gap approximately 0.35 mm (**Figure S1D**). In the middle of the hippocampus, a section of approximately 2 mm was not covered by histological slices.

Subfield delineations were based on previously described criteria in the macaque ^107^, which mirror those used in the human and previously for BigBrain ^98^. We labelled CA3 and CA4 together as one subfield because the differentiation between the two was not consistently recognizable. CA2 was evident by the high density of darkly stained neurons. The boundary between CA1 and subiculum was determined by a drop in intensities corresponding to a widening of the pyramidal cell layer. One shared label was drawn for subiculum and pre-subiculum. For a subset of slices, high-resolution 0.28 μm / pixel digital scans were available to confirm subfield definitions at higher detail. Subfield boundaries were drawn roughly orthogonal to the intrinsic spiral axis rather than oblique to have consistency across the depth of the grey matter and therefore the hippocampal surfaces. Subfield annotations were exported to geojson format to apply the TIRL registrations.

We then applied nonlinear slice-to-volume registrations to map the staining intensities from the Gallyas slices, the Cresyl Violet slices and the categorical subfield labels to the volumetric MGE space at a resolution of 0.15 mm^3^. For computational efficiency, only a hippocampal block was reconstructed in each hemisphere, rather than the whole brain. The volumetric microscopy data were then mapped to the high-resolution hippocampal surface. For representation in the flatmap space, the data were resampled to a regular matrix. To account for the gap between histological slices of the same contract, we performed linear interpolation (nearest-neighbour for subfield labels) in 2D space, which is topographically more suitable than volumetric interpolation, and smoothed the data (sigma = 3 mm). Data was sampled across all hippocampal surfaces and then averaged. The same workflow was applied to MRI metrics of microstructure following linear registration to the structural MGE volume using FSL’s FLIRT.

To account for slice-to-volume registration error and deviations in the labelling, the surface subfields map was manually corrected in GIMP v.2.8.22, guided by overlays of the microstructural surface maps as previously recommended^108^. This procedure ensured spatial continuity and plausibility of the map especially in the tail and head region of the hippocampus, where non-optimal cutting posed limitations on a serial manual labelling approach^109^. The surface mapping of the non-corrected raw subfield labels is shown in **Figure S1F**. Note that the human BigBrain subfield labels were drawn in a dense 3D volumetric reconstruction of the histological data, which meant that spatial continuity was ensured, and no such correction was needed. Applying a 3D labelling approach, however, was not applicable for the BigMac dataset, given the sparser sampling of slices and therefore the larger slice gap.

#### Subfield map quantification

We focussed quantifications of the subfield maps on the extent of the subfields along the medial-proximal axis, given the overall pattern of vertical stripes. First, we measured the global similarity of the subfield maps across hemispheres and species. A quantification of overlap (such as Dice) is not suitable for such a multilabel segmentation problem as an expansion in one subfield will affect the location of all other subfields. Therefore, we quantified the pairwise cosine-distance of each ‘row’ in the subfield map in a 4-dimensional space characterising the extent of the four subfields. The mean distance was then computed for each pair of subfield maps. Next, we quantified the relative size of each subfield compared with each other subfield. To quantify species-differences, we computed the percentage change of this metric between human and macaque.

#### Macaque *in-vivo* MRI pre-processing

In-vivo structural scans of the macaques were processed with an NHP adaptation of the openly available HCP pipeline ^97^. We adapted the pipeline scripts to run with only T1w scans as T2w scans in the same individuals were not available. Further, we adapted the brain-extraction step with an inhouse-script from the MrCat toolbox (github.com/neuroecology/MrCat). Structural processing included, amongst others, the reconstruction of the individual’s brain surface using FreeSurfer^110^ and registration to the Yerkes-19 space based on FSL’s FLIRT^111^ and FNIRT ^112^.

Volumetric processing of the resting-state fMRI scans was performed using a custom shell script pipeline of FSL commands (v.6.0, ^113^) as other automatic pipelines did not provide adequate results, particularly for brain extraction and cleaning. Initially, scans were reoriented and the five first volumes were discarded. Then the data were bias-corrected using FSL’s FAST^114^ and linearly registered with the structural scan using FLIRT. Motion correction using MCFIRT and spatial smoothing (kernel FWHM = 2 mm) was performed using FSL’s FEAT. The motion-corrected scans were further processed using ICA-based cleaning based on FSL’s MELODIC. In a first step, the non-brain-extracted scans were processed using automatic estimation of dimensionality and manually classified noise components were removed. In a second step, the cleaned and brain-extracted scans were decomposed into 20 components and the few remaining noise components were removed. The cleaned scans were then transformed to the group-level Yerkes19 template space based on non-linear registration. Following volumetric processing, the resting-state scans were processed with the fMRI surface processing part of the NHP-HCP pipeline. As part of the pipeline, the macaque fMRI data were mapped to each individual’s brain surface in fsLR_10k space and converted into cifti-format, paralleling the available human fMRI data.

All following resting-state analyses were carried out in parallel for both species using the same tools and parameters unless otherwise specified. We use the conventional term ‘cortex’ to refer to the HCP surface data, which covers mainly the neocortex. However, we do not mean to imply that the hippocampus is part of the subcortex. To assess reliability of the fMRI-based species comparison, we derived voxel-wise temporal signal-to-noise-ratio (tSNR) images for each individual and averaged these for the group. We provide the volumetric and the hippocampal surface map of tSNR in the **Supplemental Information** (**Figure S1I**).

#### Resting-state data general processing

The volumetric part of the fMRI cifti-files was mapped to the low-resolution hippocampal surface in template space (MNI for human and Yerkes19 for macaque) using the HCP’s connectome workbench toolbox (wb_command, ^115^, ribbon-constrained method). Resting-state data were then spatially smoothed on the cortical and the hippocampal surface (kernels for cortex/hippocampus: 6 mm/4 mm for the human and 2 mm/1 mm for the macaque). Prior to any further analyses, we regressed the mean time-series out of each individual’s resting-state data. To ease computational burden for gradient analyses, cortical resting-state data of each hemisphere was parcellated into approx. 1000 parcels from an existing parcellation^116^. Note that the parcels have no correspondence across species and the parcellation only served the purpose of downsampling.

#### Hippocampal gradients

The workflow for generating functional gradients was based on the Micapipe pipeline^117^ see **Figure 2A** for a schematic). In each individual, we generated a connectivity matrix between parcellated cortical and hippocampal fMRI data based on time-series correlation, followed by Fisher R-to-Z transformation. The connectivity matrices for all individuals were averaged, followed by diffusion map embedding as implemented in BrainSpace^118^. This step involved the generation of an affinity matrix using normalised angle as affinity metric. The embedding was performed for the left and right hippocampus separately, and each of them was embedded based on connectivity with cortical parcels from both hemispheres.

#### Joint cross-species embedding

Next, we generated a joint cross-species cortico-hippocampal gradient (see **Figure 2C** for a schematic). For each individual, we computed a connectivity matrix of all hippocampal vertices with all cortical parcels and averaged these for each species. Then, we concatenated the two connectivity matrices for each species so that the cortical dimension was doubled in size and applied Fisher’s transformation. Lastly, we generated an affinity matrix and applied diffusion map embedding as described above.

#### Functional cortical gradients

Cortico-cortical gradients were derived based on cross-correlation of all cortical parcels in the left and right hemispheres, followed by Fisher R-to-Z transform, affinity kernel computation, and diffusion map embedding as described above (see **Figure 3D** for a schematic). To test the relationship between cortico-cortical gradient maps and the 1^st^ joint cross-species gradient map, we performed feature selection via LASSO regression as implemented in scikit-learn (alpha = 0.1). Prior to the regression, data were transformed to a Gaussian distribution using scikit-learn’s QuantileTransformer. To compute a goodness of fit for each gradient map, we used Ordinary Least Squares regression as implemented in the Python statsmodels package. The significance of the correlations was determined using a spin test approach (1000 permutations) to control for spatial auto-correlations ^119^ as implemented in BrainSpace^118^. To quantify the overlap of gradient maps, we derived the Dice coefficient of the joint cross-species gradient and each cortical gradient map after applying a quantile-based threshold across a range of thresholds (70% – 95%). In addition, we computed the dice coefficient of the joint cross-species gradient with each pair and each triplet of cortical gradient maps.

#### Cortico-hippocampal connectivity

Connectivity of the hippocampus with the cortex was determined using Pearson correlation between each pair of vertices (**Supplemental Information, Figure S3A**). The maximal value across all hippocampal vertices was assigned to each cortical vertex to capture connectivity with any part of the hippocampus. Furthermore, we assessed whether the cortical connectivity pattern with different parts of the hippocampus is diverse (**Supplemental Information, Figure S3B**). Therefore, we constructed four hippocampal ROIs, or sectors: anterior-medial, anterior-lateral, posterior-medial, and posterior-lateral hippocampus. The four sectors were defined based on their coordinates in the hippocampal flatmap space to ensure that geometrically matched ROIs were utilised in both species. Connectivity with four hippocampal sectors was derived based on correlation with the mean time-series of each sector.

#### Homology index

Finally, we obtained a whole-brain map of species homology or divergence^47,48^, based on hippocampal connectivity (**Supplemental Information, Figure S3C**). Similar to a previous description^47,48^, we first applied a cross-species registration to establish rough correspondence of cortical parcels across species. We accessed a previously developed surface registration, which was based on cortical myelin content^94^. Then, we derived a measure of homology for each human cortical parcel: We computed the median correlation with all macaque parcels within a searchlight (radius: 15 cm) based on the spatial connectivity profile with the hippocampus. For each of the 7 human cortical networks, as defined based on the Yeo parcellation^49^, we finally computed the mean correlation.

#### Dual regression of hippocampal axes

To study the cortical reflection of the two hippocampal axes, we used a dual-regression approach ^50,51^. For each individual, cortical and hippocampal resting-state-data were concatenated in space to form a target 4D dataset. For each of the two hippocampal axes, we generated a regressor by stratifying the hippocampal flatmap (16 bins for the anterior-posterior axis and 8 bins for the proximal-distal axis) and assigning values ranging from –1 (most posterior or most proximal) to 1 (most anterior or most distal) in each bin. Those elements in the regressor corresponding to cortical and not hippocampal elements were set to 0. The two hippocampal regressors and an additional regressor modelling an intercept formed an orthogonal design matrix. In the first stage of the dual-regression we multiplied the pseudo-inverse of the design matrix with the data matrix, to obtain a single time-series for each hippocampal axis. In the second stage of the dual regression, we regressed this set of time-series back into the data matrix. This operation results in spatial brain maps quantifying the functional connectivity with each of the regressors. All individual subject’s brain maps were averaged.

#### Reprojection from cortex to hippocampal space

As alternative visualisation, we used the two group-level spatial brain maps of the dual regression analysis as coordinates for a 2-dimensional space (see **Figure 3B** for a schematic). This space reflects the intrinsic coordinates of the hippocampus itself and the aspect-ratio was adapted accordingly. The coordinates were linearly rescaled after excluding the top 10% of vertices on either end of the two axes. First, we selected a set of homologous vertices on the left hemisphere brain surface of an example subject in wb_view (**Supplemental Information, Figure S3D**) and mapped these into the hippocampal space. The cross-species difference for each region in the 2D space was assessed using Kolmogorov-Smirnov test with a significance threshold of *p* < 0.01 corrected for the number of comparisons performed. The standard deviations based on individual mappings are shown in **Supplemental Information, Figure S3E**. Next, we mapped all brain vertices of the joint cross-species gradient based on hippocampal connectivity embedding (the map shown in Figure 2C) back into this space. The two hemispheres were combined for this visualisation.

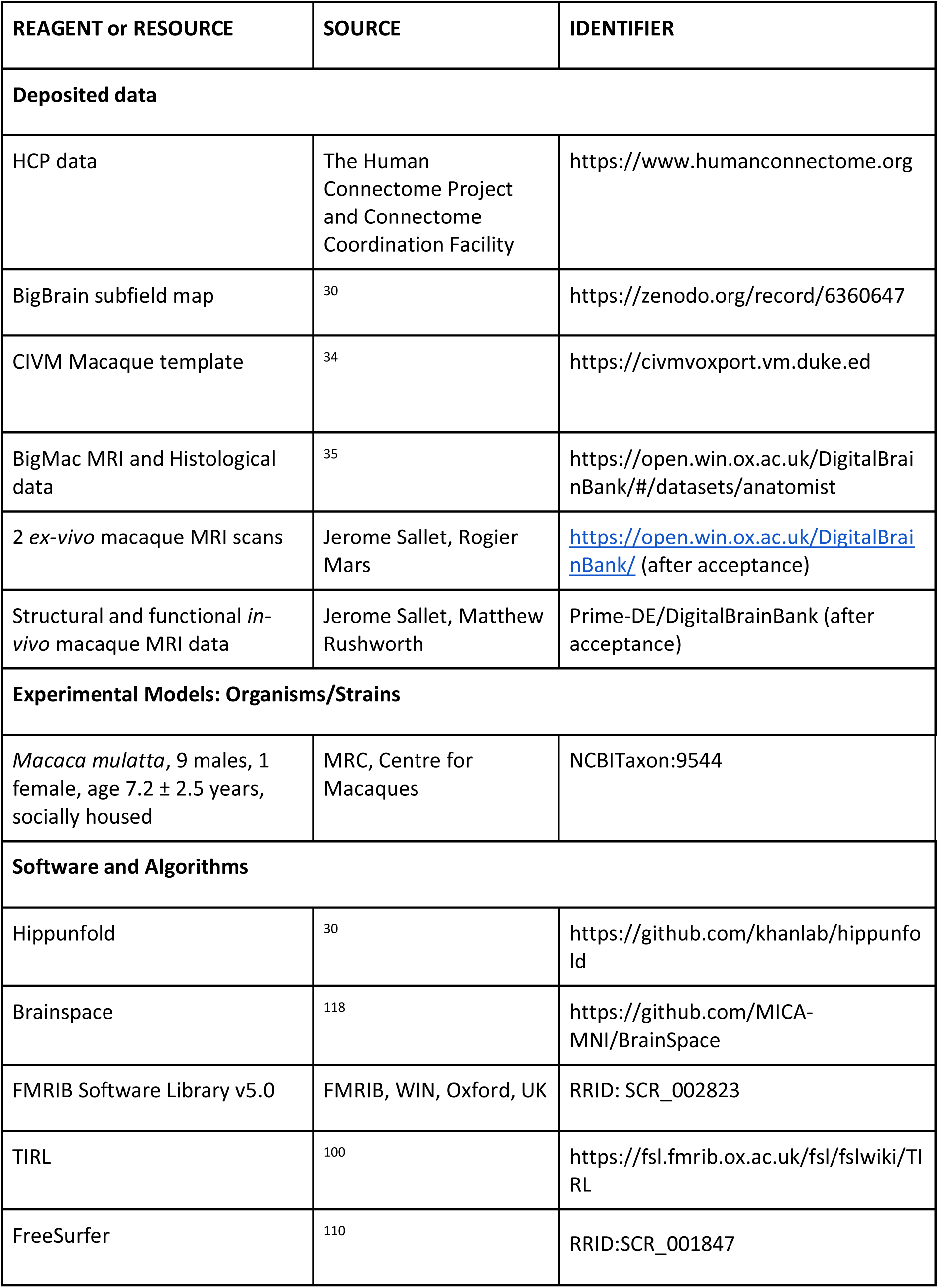
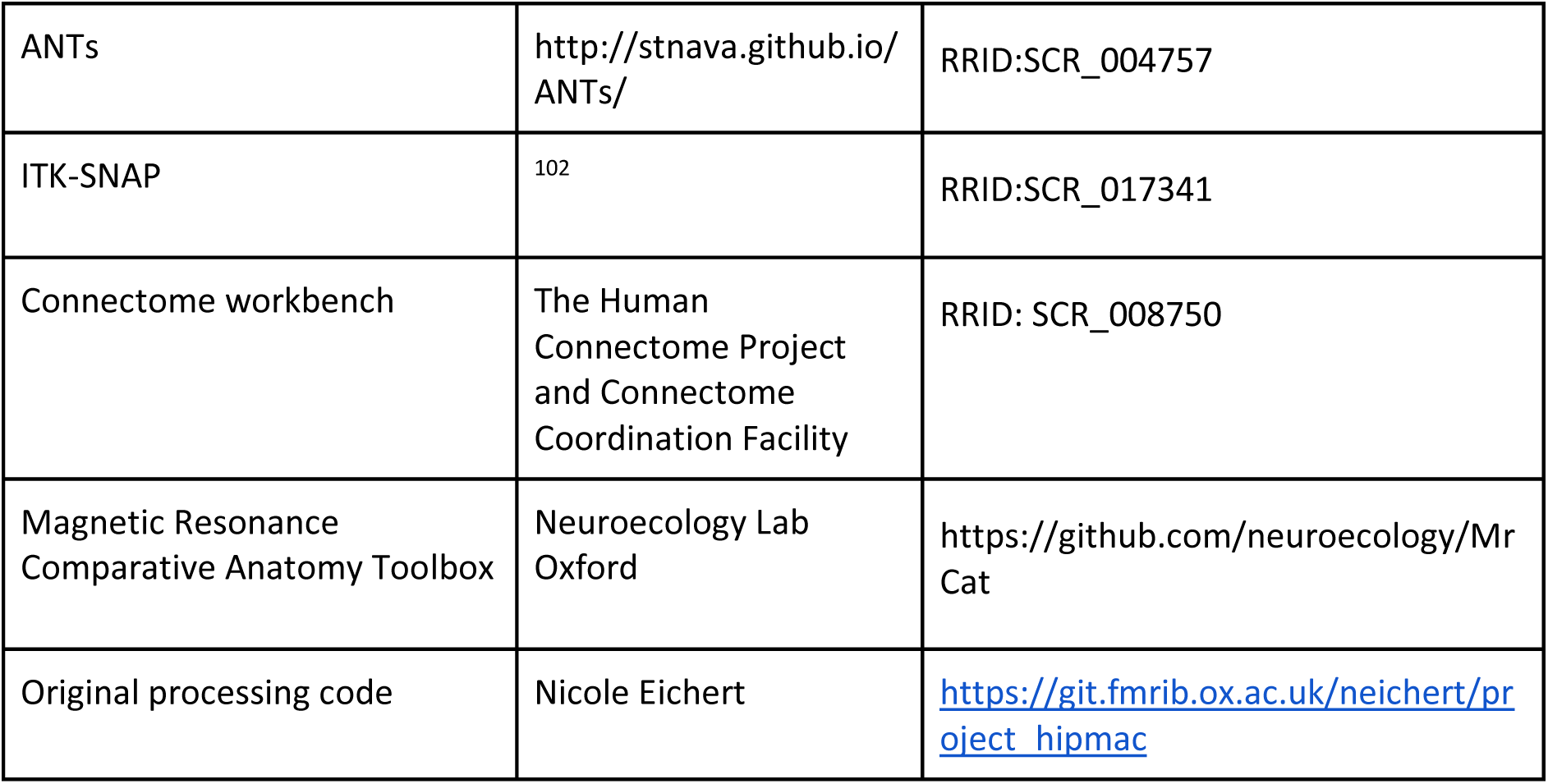
KEY RESOURCES TABLE.

### Supplemental Information

**Figure S1.**
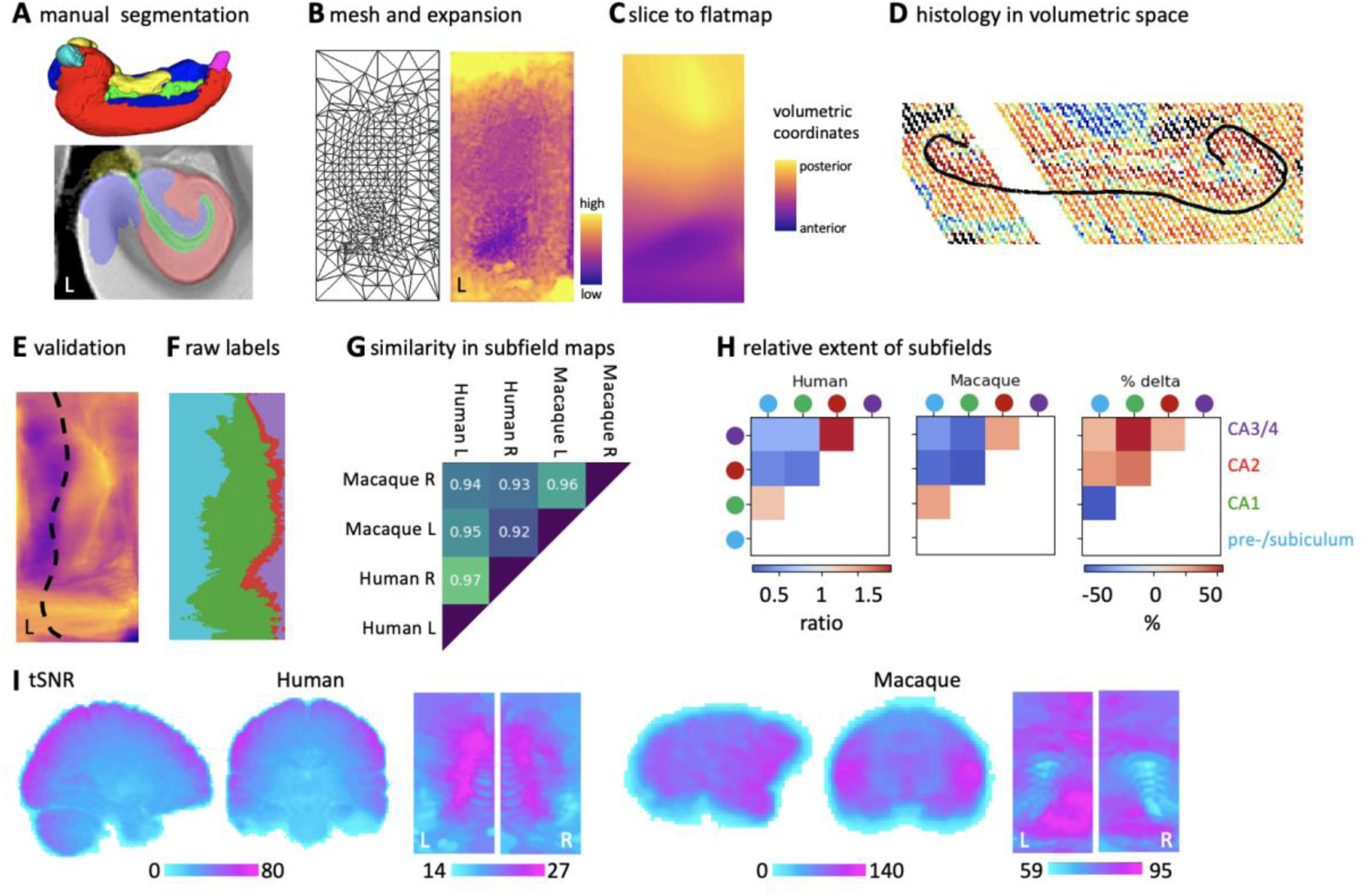
Related to Figure 1. **A**: Manual segmentations in the template brain to guide hippunfold. **B**: Low-resolution hippocampal flatmap mesh and surface area of each vertex. **C**: The volumetric order of histology slices from posterior to anterior mapped to the hippocampal flatmap. **D**: Histology mapped to 3D volumetric space and hippocampal mid-thickness surface overlaid in black. **E**: Average MGE map of two ex-vivo macaque brains. Dashed line: CA1-subiculum boundary from BigMac. **F**: Raw subfield labels mapped to the surface before correction on the surface. **G**: Cosine similarity of the hippocampal subfield maps across hemispheres and species. **H**: Extent of subfields relative to each other in human and macaque and percentage difference between human and macaque. **I**: Average tSNR maps (*n* = 10) of the rs-fMRI data for human and macaque shown in volumetric space and sampled to the hippocampal surface.

**Figure S2.**
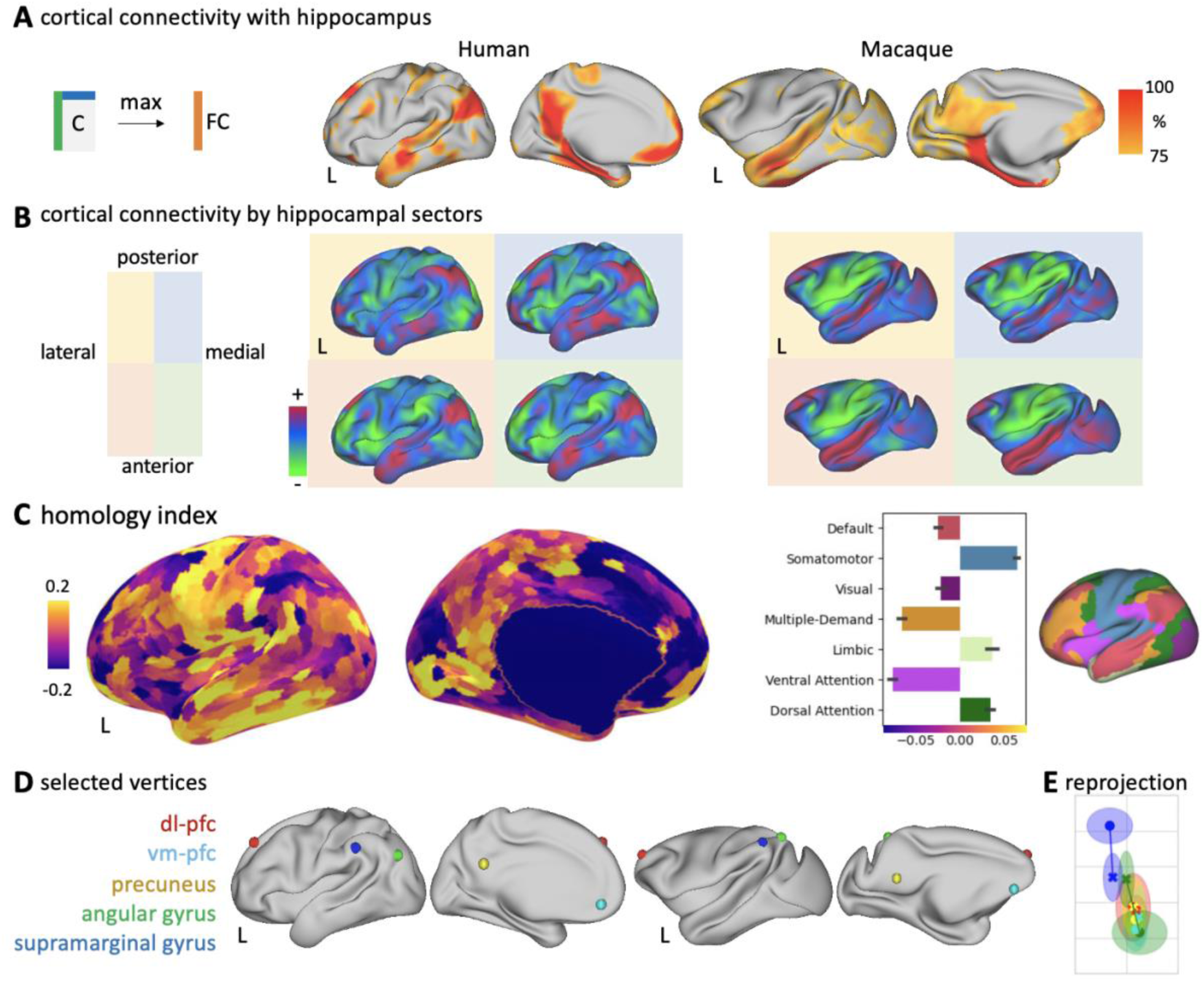
Related to Figure 3. **A**: Left – In each species, a functional connectivity map (FC) was obtained by computing the maximal connectivity of each cortical vertex from the cortico-hippocampal connectivity matrix (C). Right – Thresholded functional connectivity map (>75%, *n*=10). **B**: Functional connectivity with average signal from four spatial sectors of the hippocampus. **C**: Homology index computed based on hippocampal connectivity profiles after applying a cross-species registration. Left – Whole-brain map. Right – Mean homology index per network^49^. **D**: Selected vertices mapped in Figure 3B. **E**: Mean and standard deviation of the coordinates plotted in Figure 3B based on 10 individual subjects. Note the mean location in the 2D space is slightly different than in Figure 3B because the quantile-based thresholding of the axis in individuals differs from the threshold based on the group average.

**Figure S3.**
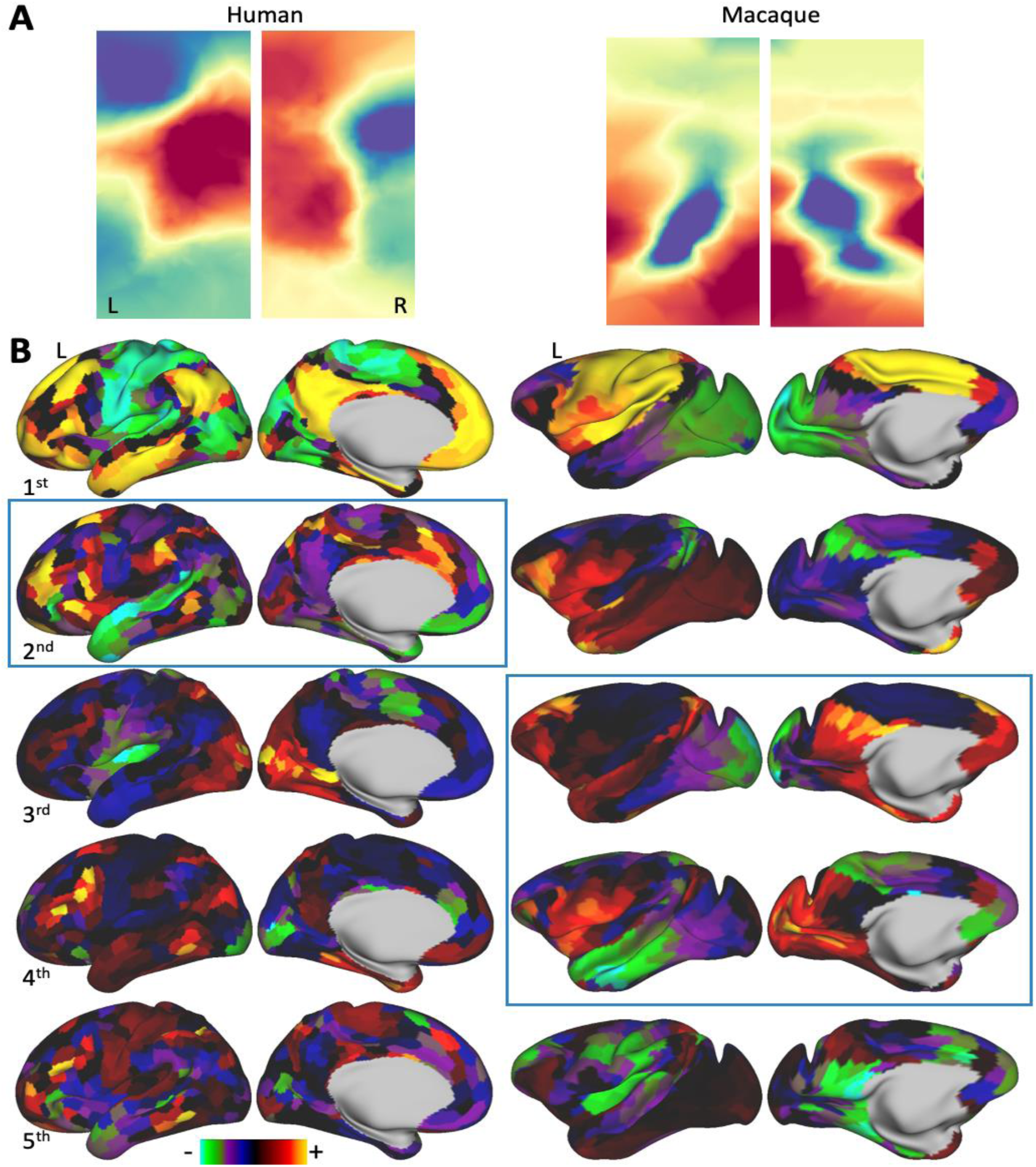
Related to Figure 2. **A**: Higher-order hippocampal gradients: 2^nd^ human and 6^th^ macaque hippocampal gradient displaying proximal-distal differentiation. The sign of the map is random. **B**: Cortico-cortical gradients. The boxes highlight the gradients that match the joint cross-species gradient, i.e., the thresholded maps shown in Figure 2D.

